# Multiplex secretome engineering enhances recombinant protein production and purity

**DOI:** 10.1101/647214

**Authors:** Stefan Kol, Daniel Ley, Tune Wulff, Marianne Decker, Johnny Arnsdorf, Jahir M. Gutierrez, Austin W.T. Chiang, Lasse Ebdrup Pedersen, Helene Faustrup Kildegaard, Gyun Min Lee, Nathan E. Lewis

## Abstract

Host cell proteins (HCPs) are process-related impurities generated during biotherapeutic protein production. HCPs can be problematic if they pose a significant metabolic demand, degrade product quality, or contaminate the final product. Here, we present an effort to create a “clean” Chinese hamster ovary (CHO) cell by disrupting multiple genes to eliminate HCPs. Using a model of CHO cell protein secretion, we predicted the elimination of unnecessary HCPs could have a non-negligible impact on protein production. We analyzed the total HCP content of 6-protein, 11-protein, and 14-protein knockout clones and characterized their growth in shake flasks and bioreactors. These cell lines exhibited a substantial reduction in total HCP content (40%-70%). We also observed higher productivity and improved growth characteristics, in specific clones. With the reduced HCP content, protein A and ion exchange chromatography more efficiently purified a monoclonal antibody (mAb) produced in these cells during a three-step purification process. Thus, substantial improvements can be made in protein titer and purity through large-scale HCP deletion, providing an avenue to increased quality and affordability of high-value biopharmaceuticals.

## Introduction

Host cell proteins (HCPs) that are released from dead cells and secreted from viable cells accumulate extracellularly during mammalian cell culture, potentially impairing product quality^1,2^ and posing an immunogenic risk factor^3^. HCPs must be reduced to low levels (less than 1–100 ppm) in all cell-derived protein biotherapeutics before final product formulation, putting a great demand on downstream purification. Depending on titer, downstream processing comprises up to 80% of the entire production costs of a monoclonal antibody (mAb)^4^. In the production of a mAb, protein A affinity capture is often employed as a generic first step, followed by two or three orthogonal polishing steps^5^. Although the concentrations of HCPs in cell culture harvests are reduced to acceptable levels after these purification steps, certain HCPs can escape an entire purification process and remain in the final product at levels that may affect product quality and stability, cause an immunogenic reaction, or display residual activity^6,7^. It is therefore of utmost importance to closely monitor and identify HCPs to ensure their removal from biotherapeutic products. Enzyme-linked immunosorbent assay (ELISA) is typically used in industrial processes to monitor the total HCP content, but gives no information on specific HCP components^8^. Proteomic analysis, such as two-dimensional gel electrophoresis and/or mass spectrometry (MS), can therefore be employed to identify specific HCPs^6,9,10^. Although many studies have focused on identification, only a few efforts have been made to remove troublesome HCP moieties by host cell line engineering^1,2,11,12^.

Concerning CHO HCPs that affect product quality, a matripase-1 knockout cell line was recently created that prevents degradation of recombinant proteins^2^. A knockout cell line has also been created devoid of lipoprotein lipase (Lpl) activity^1^. Lpl is a difficult-to-remove HCP that degrades polysorbate in mAb formulations, and targeted mutation resulted in abolishment of Lpl expression and reduced polysorbate degradation without substantial impact on cell viability. In a recent study, the HCPs Annexin A2 and cathepsin D were removed^12^. Although the authors do not confirm loss of activity, the knockouts do not adversely affect cell growth. In addition, knockout of a serine protease was shown to eliminate proteolytic activity against a recombinantly expressed viral protein^11^. These previous efforts targeted single genes, but mammalian cells can secrete a few thousand HCPs^9^, so CRISPR technology could be used to target multiple genes simultaneously by simply expressing multiple sgRNAs together with a single Cas9 nuclease^13,14^.

Here, we demonstrate substantial reductions in HCP content can be obtained through targeted genome editing, guided by omics analyses. Specifically, we used a systems biology computational model to show that the removal of multiple HCPs could free up a non-negligible amount of energy. We then used proteomics to help identify target HCPs and subsequently applied multiplex CRISPR/Cas9 to disrupt up to 14 genes coding for proteins that are abundant in harvested cell culture fluid (HCCF), difficult-to-remove during downstream processing, or have a potentially negative impact on product quality. We analyzed HCP content of a 6-protein, 11-protein, and 14-protein knockout and characterized their growth in shake flasks, Ambr bioreactors, and DASGIP bioreactors. We observed a substantial reduction of total HCP content in the 11×KO (∼40%), 11×KO and 11×KO (60-70%) cell lines. Depending on the cell line and growth conditions, we also observed increased mAb productivity and improved growth characteristics. When measuring HCP content and mAb titer at the different stages of a three-step purification process (using protein A and two ion exchange chromatography steps), we observed a strong reduction in HCP content, increased HCP log reduction values (LRV) of protein A affinity chromatography, and decreased HCP ppm values. Thus, we demonstrate that superior CHO production host cell lines can be produced by eliminating specific HCPs. The dramatically lowered HCP ppm values during purification of a mAb will facilitate downstream processes. These HCP-reduced knockouts can be combined with additional advantageous genetic modifications, such as the production of defucosylated antibodies^15^, higher viability during long culture times^16^, and/or clones with higher protein production stability of mAbs^17^, to create predictable upstream and downstream processes with full control over critical process parameters.

## Materials and Methods

### Cell culture and cell line generation

All cell lines described here were created using CHO-S cells (Life Technologies, Carlsbad, CA, USA). Knockout cell lines were generated as previously described^14^. Viability, cell size and VCD was monitored using the NucleoCounter NC-200 Cell Counter (ChemoMetec, Allerod, Denmark). Batch cultivation was performed in 250 mL shake flasks containing 60-80 mL CD-CHO medium (Thermo Fisher Scientific, Waltham, MA, USA) supplemented with 8 mM L-glutamine (Life Technologies) and 0.2% anti-clumping reagent (Life Technologies). Cells were extracted from the cell bank and thawed in 10 mL pre-warmed medium. Cultures were passaged three times during 7 days and strained through a 40 µm filter before inoculation. Cultivation was performed in a humidified shaking incubator operated at 37°C, 5% CO_2_ and 120 rpm.

Prior to inoculation into fed-batch bioreactors, cells were adapted from CD-CHO medium to FortiCHO (Gibco, Carlsbad, CA, USA), OptiCHO (Gibco), or ActiPro (GE Healthcare, Chicago, IL, USA) medium during three passages. All media were supplemented with 1% Anti-Anti (Life Technologies), 0.1% anti-clumping agent (Life Technologies) and 8 mM L-glutamine (Lonza, Basel, Switzerland). Fed-batch cultivation was performed in a DASGIP Mini bioreactor (Eppendorf, Jülich, Germany) containing between 260-280 mL medium or in an Ambr 15 bioreactor system (Sartorius, Göttingen, Germany) containing 14.4 mL medium. Temperature was maintained at 37°C. In the DASGIP, dissolved oxygen (DO) was maintained at 40% by cascade aeration strategy (Air/Air&O_2_/O_2_) using nitrogen as a carrier gas (total flow rate 1-3 L/h). In the Ambr, DO was maintained at 40% using pure O_2_ using nitrogen as carrier gas (total flow rate 0.200 sL/h). Agitation rate (200-400 rpm) was adjusted on a daily basis to keep DO set point at 40%. Culture pH was maintained at 7.1 ±0.02 (DASGIP) or 7.1 ± 0.05 (Ambr) using intermittent addition of CO_2_ to the gas mix and 1 M CHNaO_3_. HyClone Cell Boost 7a supplement (GE Healthcare) containing 500 mM glucose and HyClone Cell Boost 7b Supplement (GE Healthcare) were fed daily when cell numbers reached 1.4 − 2.2 × 10^6^ cells/mL at 2% and 0.2% of daily total volume, respectively. Glucose (2220 mM) and glutamine (200 mM) were used to maintain levels between 24-30 mM and 3-5 mM, respectively, via a once-daily feeding based on measured concentration, doubling time and consumption rates. Anti-foam C (Sigma, St. Louis, MI, USA) was supplemented when needed.

### Deep sequencing analysis of genome edit sites

Confluent colonies from 96-well flat-bottom replicate plates were harvested for genomic DNA extraction. DNA extraction was performed using QuickExtract DNA extraction solution (Epicentre, Illumina, Madison, WI) according to the manufacturer’s instruction. The library preparation was based on Illumina 16S Metagenomic Sequencing Library. Preparation and deep sequencing were carried out on a MiSeq Benchtop Sequencer (Illumina, San Diego, CA). The protocol for amplifying the CRISPR-targeted genomic sequences, amplicon purification, adapter-PCR and quality analysis followed our previously published methods^14^.

### Generation of antibody expressing cell lines

WT and knockout cell lines (11×KO, 11×KO and 11×KO) were initiated from cryopreserved vials and passaged for one week prior to transfection. Transfection efficiency was determined by transfecting the pMAX-GFP (Lonza) and subsequent FACS analysis. Cells were transfected with a plasmid encoding the antibody Rituximab and the zeocin selection gene using Freestyle MAX reagent, following the manufacturer’s recommendations and maintained for two days before selection pressure was initiated with zeocin (400 µg/mL). After 21 days of selection, viability was above 95% in all cultures and pools were stored in vials containing 10% DMSO. To enrich for clones with high productivity, immunostaining of Rituximab on the plasma membrane (surface staining) was performed on the Rituximab expressing cell pool by staining with an anti-human IgG antibody conjugated to FITC (Invitrogen, Carlsbad, CA, USA) and subsequent single-cell sorting using fluorescence activated cell sorting (FACS) and expansion. To select clones with the highest productivity, we used the titer-to-confluency method^18^. Confluency was determined on a Celigo cytometer (Nexcelom Bioscience, Lawrence, MA, USA) and titer was determined using biolayer interferometry. The number of clones screened by this method was 1000 (WT), 480 (11×KO), 1760 (11×KO), and 480 (11×KO).

### Quantification of secretome costs

A genome-scale model of protein secretion was adapted for use here^19^. Briefly, this model was adapted to focus on the bioenergetic costs of protein synthesis and translocation of each host-cell secreted protein. Information on each secreted protein in CHO cells was obtained, including mRNA and protein sequence, signal peptides, etc. We then simulated the cost of producing each protein. Each cost was scaled by the measured mRNA levels using published RNA-Seq from CHO-S cells grown under similar conditions^20^, and this was used as a proxy for the relative resources allocated to each secreted protein. Entrez gene identifiers were matched to their corresponding entry number in the UniProt database to determine presence of signal peptide. For those genes without a UniProt match, we used the online tool PrediSi^21^ to determine the presence of a signal peptide. To estimate ATP cost of secretion for all secreted genes identified, we added the following costs, adapted from the genome-scale model of protein secretion^19^. First, the energy cost of protein translation was equal to 4 × L ATP molecules where L is the length of the amino acid sequence. Next, the average cost of signal peptide degradation was equal to 22 ATP molecules. Finally, the energetic cost of translocation across ER membrane was equal to L/40 + 2. From this, we were able to quantify what proportion of all secreted protein was eliminated in our study.

### Mass spectrometry identification of HCPs

WT CHO-S was cultivated in duplicate in three separate DASGIP Mini bioreactor experiments. HCCF was obtained during late exponential phase by centrifugation at 500*g* for 10 minutes. Samples were TCA-precipitated with an overnight incubation in ice cold acetone, and the dry protein pellet was dissolved in 100 µl 8M urea. Samples were treated first with 100 mM DTT (5 µl) and incubated at 37°C for 45 minutes and then with 100 mM iodoacetamide (10 µl) and incubated in the dark for 45 minutes. Samples containing 100µg of protein were digested overnight at 37°C using trypsin after which 10% TFA was added, and samples were stage tipped. Data were acquired on Synapt G2 (Waters) Q-TOF instrument operated in positive mode using ESI with a NanoLock-spray source. During MS^E^ acquisition, the mass spectrometer alternated between low and high energy mode using a scan time of 0.8 s for each mode over a 50-2000 Da interval. Nanoscale LC separation of the tryptic digested samples was performed using a nanoAcquity system (Waters, Milford, MA, USA) equipped a nanoAcquity BEH130 C18 1.7 µm, 75 µm × 250 mm analytical reversed-phase column (Waters). A reversed-phase gradient was employed to separate peptides using 5–40% acetonitrile in water over 90 min with a flow rate of 250 nL/min and a column temperature of 35°C. The data were analyzed using the progenesis QI software (NonLinear dynamics), which aligns the different runs ensuring that the precursor ions and identifications can be shared in between runs.

### HCP, antibody and protein quantification

All quantification was performed using biolayer interferometry on an Octet RED96 (Pall, Menlo Park, CA, USA). For HCP quantification, the anti-CHO HCP Detection Kit (Pall) was used according to the manufacturer’s specifications. When needed, samples were diluted before analysis. For Rituximab quantification, protein A biosensors (Pall) were equilibrated in sample diluent buffer (PBS containing 0.1% BSA and 0.02% Tween-20), and Rituximab was measured for 120s at 30°C. Absolute concentrations of Rituximab were calculated by comparison with a calibration curve generated from a dilution series of human IgG (A01006, Genscript, New Jersey, NJ, USA). Regeneration of biosensor tips between measurements was performed in 10 mM glycine pH 1.7. When needed, samples were diluted to fall within the range of the calibration curve. The protein content of HCCF and cells was measured using a Nanodrop 2000 (Thermo Fisher Scientific). Before analysis, HCCF was TCA-precipitated, and whole cells were lysed using RIPA buffer.

### Protein purification

Cell-free supernatants were prepared by centrifugation and directly loaded (100 mL) onto a 5 mL HiTrap MabSelect column (GE Healthcare). After washing the column with binding buffer (20 mM sodium phosphate, 0.15 M NaCl, pH 7.2), Rituximab was eluted with elution buffer (0.1 M sodium citrate, pH 3.5) into collection tubes containing 1 M TRIS-HCl, pH9.0. The fractions containing Rituximab were pooled, diluted 10 times with 50 mM MES pH 6.0, and loaded onto a 5 mL HiTrap SP FF column (GE Healthcare). After washing the column with binding buffer (50 mM MES pH 6.0, 40 mM NaCl), Rituximab was eluted with elution buffer (50 mM MES pH 6.0, 200 mM NaCl). The fractions containing Rituximab were pooled, diluted 5 times with 20 mM TRIS pH 8.0, and loaded onto a 5 mL HiTrap Q FF column (GE Healthcare) equilibrated in buffer A (20 mM TRIS-HCl, 40 mM NaCl). The flow through containing Rituximab was collected and concentrated to 5 mg/mL. The final mAb samples were aliquoted, snap frozen in N_2_ (l) and stored at −80°C. Samples for analysis were collected after every chromatographic step.

## Results

### CHO cells spend substantial resources on the host cell secretome

There are several potential advantages to the removal of HCPs, including the elimination of immunogenic proteins or proteins that degrade product quality. However, the question remains regarding the bioenergetic investment made by the host cells on the secreted fraction of their proteome. To evaluate the resource and bioenergetic investment of CHO cells into their secretome, we calculated the cost of each protein using a model of CHO cell protein secretion^19^, scaled by its expression in RNA-Seq data^20^. In CHO-S cells, we found that 39.2% of the resources are dedicated to producing membrane and secreted proteins, most of which are processed through the secretory pathway (Fig. 1A). 10.2% of the cell resources were explicitly dedicated to secreted proteins with signal peptides, and a relatively small number of secreted proteins account for the majority of the bioenergetic cost and use of the secretory pathway. Furthermore, we anticipate that the cost for secreted proteins is likely higher since these proteins are not turned over and recycled like intracellular and membrane proteins. Thus, there is great potential to free up cellular resources and secretory capacity, in addition to improving product quality by eliminating unnecessary HCPs.

**Figure 1.**
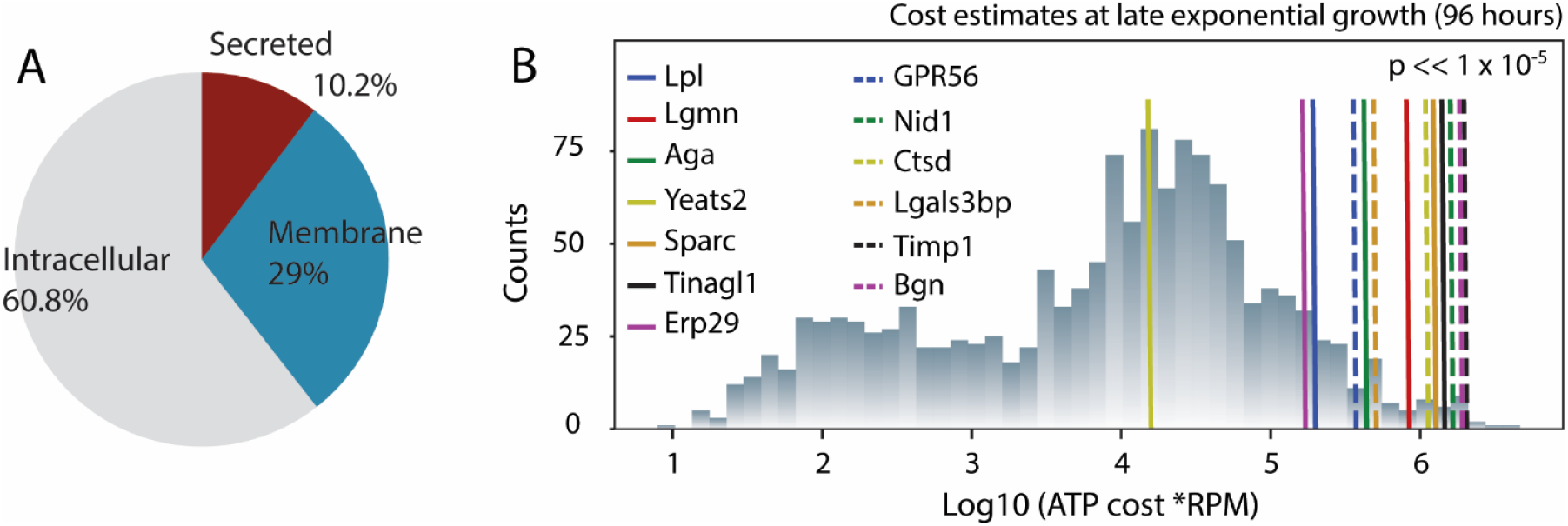
Substantial cellular resources can be liberated by targeting HCPs in the CHO secretome. (A) Proteins produced through the secretory pathway account for more than 40% of the cell’s biosynthetic capacity, and roughly 1/4 of this cost is dedicated to secreted proteins. (B) Following proteomic analysis of the HCCF, 14 proteins (associated with 13 gene loci) were identified and targeted for deletion. We quantified the bioenergetic cost of protein synthesis and secretion in CHO cells, and we found the deletion of the target genes will free up significantly more bioenergetic resources than expected from the deletion of a similar number of randomly selected genes. RPM = reads mapping to gene per million reads in the transcriptome.

### Target selection and verification

To decrease HCP secretion and free up resources for CHO cell growth and protein production, we identified targets based on three criteria: (i) being abundant in CHO supernatants as analyzed by mass spectrometry (Table S1), (ii) being a difficult-to-remove impurity in downstream processing^22–25^, and (iii) having a negative impact on product quality^1,26,27^. To remove the genes, we subjected the cell lines in this study to 4 cycles of CRISPR/Cas9-mediated multiplex gene disruption. In the first round, the gene encoding Metalloproteinase inhibitor 1 (Timp1) was disrupted. In the second round, the genes encoding Biglycan (BGN), Galectin-3-binding protein (LGALS3BP), Nidogen-1 (NID1.1 and NID1.2), and Cathepsin D (CTSD) were disrupted, resulting in the 11×KO cell line. In the third round, the genes encoding N(4)-(Beta-N-acetylglucosaminyl)-L-asparaginase (Aga), Endoplasmic reticulum resident protein 29 (Erp29), G-protein coupled receptor 56 (Gpr56), Tubulointerstitial nephritis antigen-like (Tinagl1), and Legumain (Lgmn) were disrupted, resulting in the 11×KO cell line. Finally, in the fourth round, we disrupted the genes encoding YEATS domain-containing protein 2 (Yeats2), SPARC (Sparc), and Lipoprotein lipase (Lpl), resulting in the 11×KO cell line.

The targets were predominantly identified through proteomics on spent media or purified product, and we verified that these genes were, on average among the more abundant transcripts for secreted proteins, and accounted for proteins exerting a greater cost to the cell (Fig. 1B). Furthermore, upon quantifying the amount of ATP liberated upon their deletion, we found their deletion led to a significantly higher amount of resources eliminated (p << 1×10^−5^), when compared to a comparable number of randomly selected genes.

We verified gene disruption at the genomic and proteomic levels. Specifically, we used MiSeq analysis (Table 1) to verify that all indels led to frameshift mutations except for the one generated in LGALS3BP, which leads to a partly scrambled amino acid sequence and the introduction of a stop codon after residue 44. Mass spectrometry was subsequently used to verify the absence of our target proteins after gene disruption. Supernatants from wild-type CHO-S (WT) and the 11×KO, 11×KO and 11×KO cell lines were analyzed for the presence of peptides derived from the target proteins (Fig. 2). Except for the target YEATS2, peptides were detected for all of the targets in WT. No or very few peptides were detected in the 11×KO, 11×KO and 11×KO cell lines as compared to WT for all targets except LGALS3BP. Approximately 10-20% of LGAL3SBP peptides derived from the region upstream of the editing site were still present in all knockout cell lines, indicating secretory down-regulation of the partial non-sense protein produced by the frameshift mutation. All other targets were no longer detected.

**Table 1.**
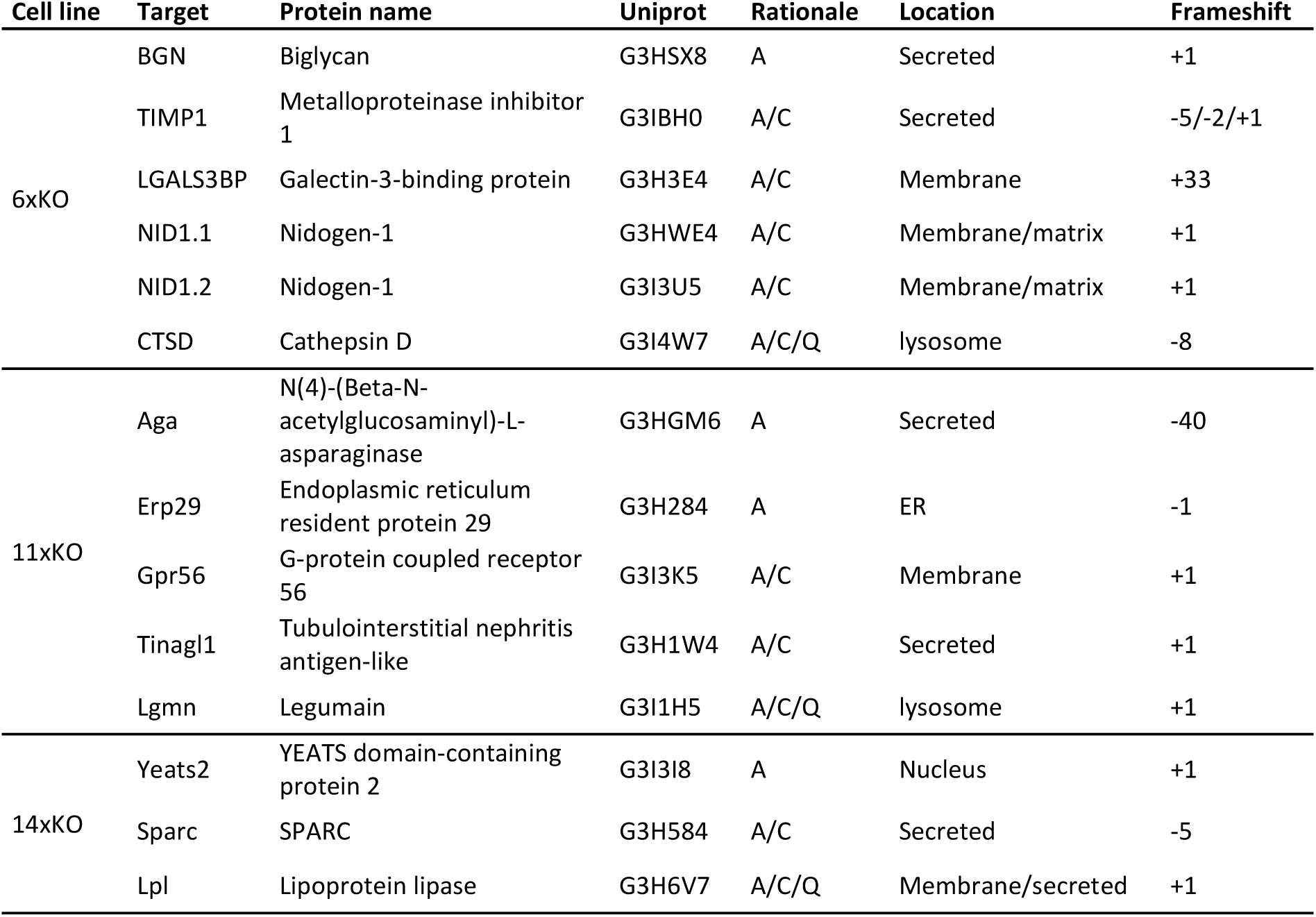
Selection and MiSeq verification of knockout targets addressed in this study. The target genes names, protein names, UniProt identifiers, rationale for deletion, subcellular location, and indel size are indicated. For rationale: A, abundance; C, copurifying; Q, quality.

**Figure 2.**
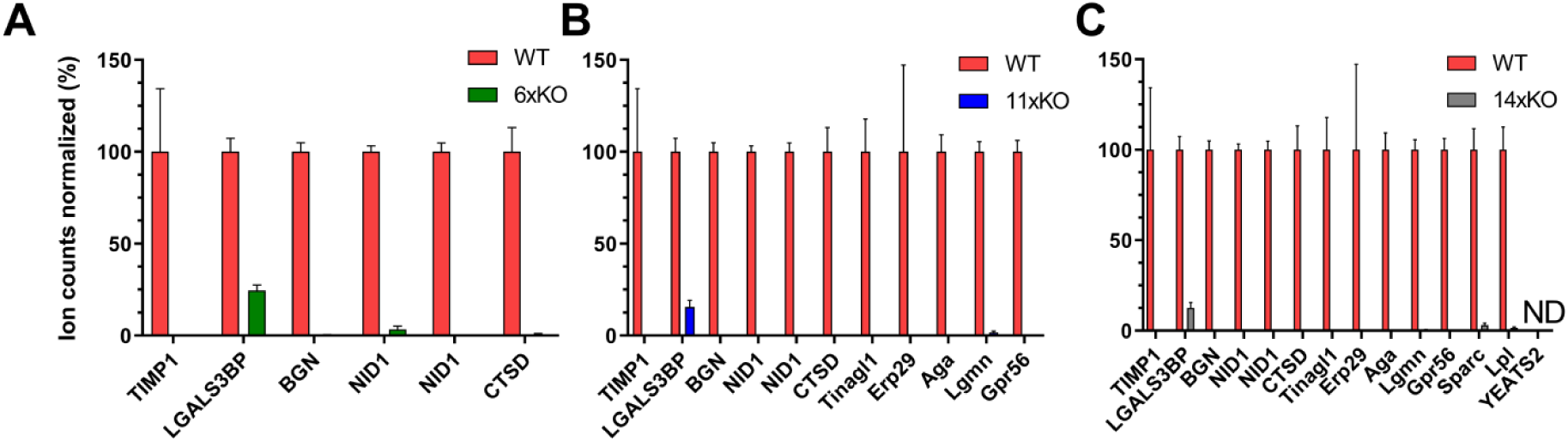
Verification of knockouts by mass spectrometry. Supernatant samples prepared from WT (red bars) were subjected to MS and compared to the 11×KO (A; green bars), 11×KO (B; blue bars) and 11×KO (C; grey bars) cell lines with respect to the indicated targets. Ion counts were normalized to the levels detected in WT. The target YEATS2 was not detected (ND) in both WT and the 11×KO cell line. Mean and standard deviation of three technical replicates are shown.

### Knockout cell lines display improved growth characteristics and decreased HCP content in batch cultivation

To assess the properties of our knockout cell lines, we followed their growth and viability for 7 days in shake flasks with daily sampling to measure viable cell density (VCD), cell viability, and HCP content (Fig. 3). Interestingly, cell density and viability of knockout cell lines were improved over WT (Fig. 3A). While the 11×KO cell line displayed an intermediate phenotype, growth of the 11×KO and 14×KO cell lines was enhanced to reach a VCD of approximately 7.6 million cells per milliliter. WT reached a VCD of approximately 5.4 million cells per milliliter. Viability of WT started to decrease on the fifth day of culture, while the knockout cell lines remained above 90% viability for the duration of the experiment. Samples were analyzed using a CHO HCP detection kit and showed a remarkable reduction in HCP content as compared to WT (Fig. 3B). Whereas the 11×KO cell line shows an intermediate phenotype again, the 11×KO and 11×KO cell lines behave similarly, showing an HCP reduction to approximately 150 µg/mL. As the VCD of the knockout cell lines is higher, the behavior of the cell lines can arguably better be described by calculating specific HCP productivities. Specific HCP productivity was found to be 13.5 picograms per cell per day (pcd) for WT, whereas the 11×KO, 11×KO and 11×KO cell lines were reduced to 9.9, 5.7 and 5.6 pcd, respectively (Fig. 3C). Specific HCP productivity was found to be reduced by 27%, 58% and 59% for the 11×KO, 11×KO and 11×KO cell lines, respectively, as compared to WT (Fig. 3D). Besides the improvements in growth characteristics during batch cultivation, we encountered an unexpectedly high reduction in HCP content. As it could be argued that the HCP-reduced phenotype is an artifact of the knockout procedure, we determined the HCP content of unrelated cell lines harbouring multiple knockouts generated using the same procedure. No significant decrease in HCP content of WT CHO-S and the knockout cell lines is observed (Fig. S1 and accompanying text), showing that the HCP-reduced phenotype is specifically caused by the removal of our targeted proteins. To gain a better understanding of the impact of these knockouts, we measured several basic cell properties and established whether mAb productivity was affected.

**Figure 3.**
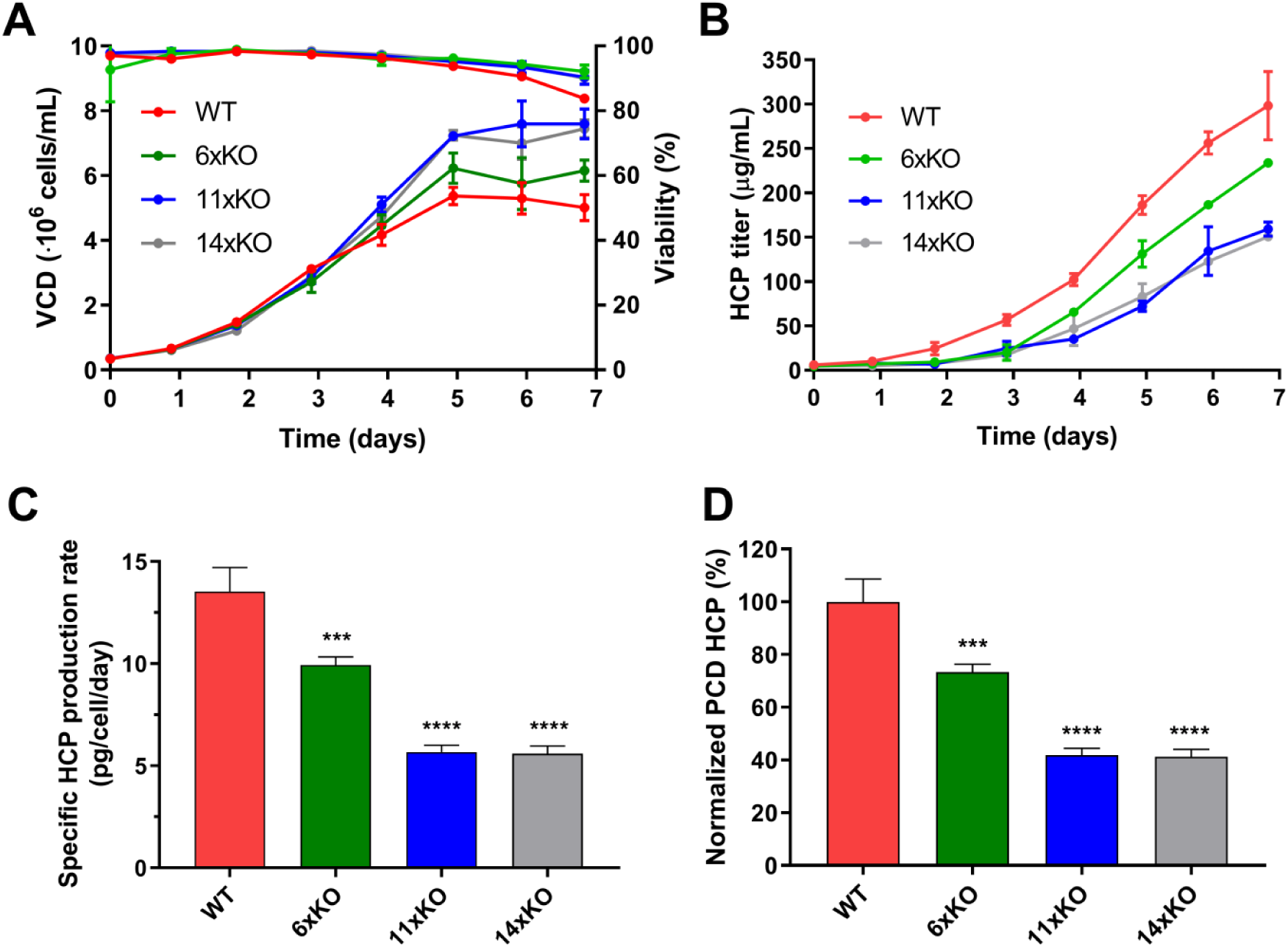
Knockout cell lines display improved growth properties and HCP profile. Viability and VCD (A), HCP profile (B), normalized HCP content on day 7 (C), and normalized specific HCP productivity (D) of WT (red), 11×KO (green), 11×KO (blue), and 11×KO (grey) cell lines. Viability, VCD and HCP content were measured from day 0 to day 7 in shake flasks. The specific HCP productivity was determined as the slope of HCP concentration versus the integral number of viable cell (IVCD) calculated from day 0 to day 7, and expressed as pg HCP per cell and per day (HCP pcd). Specific HCP production was normalized to the levels detected in WT. Mean and standard deviation of three technical replicates are shown. Statistical analysis was performed using one-way ANOVA followed by Tukey’s post-hoc test (*** P ≤ 0.001, **** P ≤ 0.0001).

### Antibody-producing knockout cell lines display higher productivity and less total HCCF protein content

To assess several basic cell properties and protein productivity, we generated stable pools expressing the monoclonal antibody Rituximab (Rit). We cultivated the WT, WT Rit, 11×KO Rit, 11×KO Rit, and 11×KO Rit cell lines for 4 days in shake flasks and collected HCCF and cell pellets. We analyzed HCP content, protein content of HCCF and cells, cell size, transfection efficiency, antibody staining efficiency, and Rituximab titer (Fig. 4 and Fig. S2). The HCP-reduced phenotype was observed as before (Fig. S2B), while VCD was similar for all cell lines (Fig. S2A). The total protein content of the HCCF obtained from the knockout cell lines was significantly reduced by approximately 50% (Fig. 4A), while cell size was reduced from 13.4 µm to 12.5 µm (Fig. S2B). The protein content per cell remained unchanged at approximately 150 pg protein per cell (Fig. 4B), which corresponds well with published values using a CHO-K1 cell line^28^. Antibody productivity in cell pools was assayed in two ways: 1) by an antibody staining method and subsequent FACS analysis (Fig. 4C) and 2) by biolayer interferometry (Fig. 4D). The knockout cell lines displayed higher productivity as compared to WT using both methods. Whereas productivity increased with the number of knockouts using the staining method, it is only increased until the 11×KO cell line using biolayer interferometry. Transfection efficiency was similar for all cell lines (Fig. S2D). Based on these results, we selected high-producing clones to analyze their growth properties under fed-batch conditions. During selection, the 11×KO Rit cell line displayed a low single cell survival rate and no clones could be selected that displayed sufficiently high mAb productivity. Based on these observations and as the HCP-reduced phenotype was not decreased further in the 11×KO cell line, we excluded the 11×KO Rit cell line from further study.

**Figure 4.**
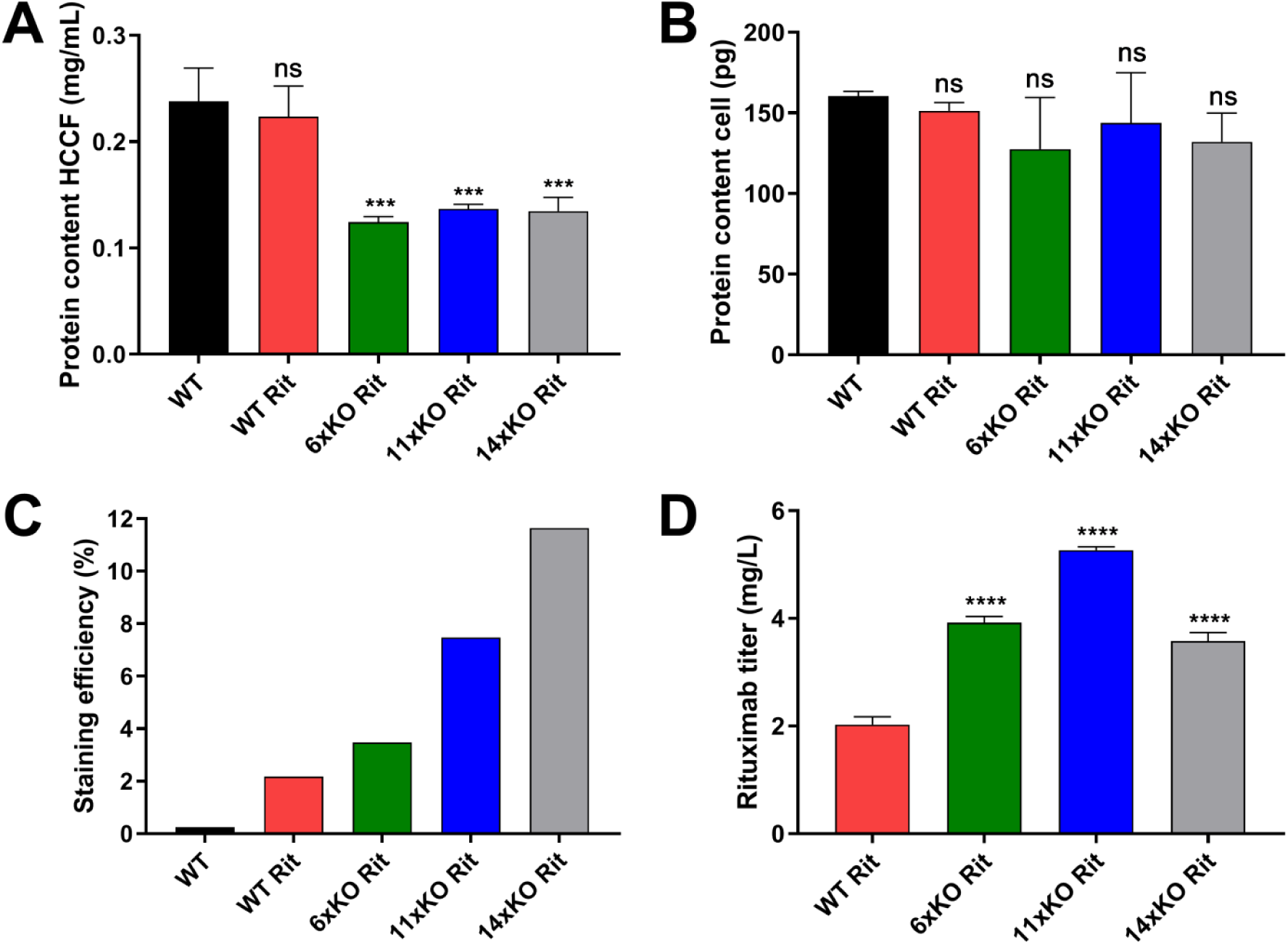
Antibody-producing pools display increased productivity and less HCCF protein content. HCCF protein content (A), protein content per cell (B), staining efficiency (C), and Rituximab titer (E) of WT (black), WT Rit (red), 11×KO Rit (green), 11×KO Rit (blue), and 11×KO Rit (grey) cell lines. All properties were measured after cultivation for four days in shake flasks. Mean and standard deviation of three technical replicates are shown for all experiments except for the staining efficiency determination. Statistical analysis was performed using one-way ANOVA followed by Tukey’s post-hoc test (ns = not significant, *** P ≤ 0.001, **** P ≤ 0.0001).

### Antibody-producing KO clones display decreased HCP content in fed-batch cultivation

After selecting high mAb-producing clones of the WT Rit, 11×KO Rit, and 11×KO Rit cell lines, we analyzed their growth properties, productivity and HCP content in Ambr (Fig. S3) and DASGIP bioreactors (Fig. 5). The goal of these experiments was to compare behavior of clones in different media and to generate material for downstream processing. In addition, we wanted to assure that high producers could be generated using the knockout cell lines and that these retained the HCP-reduced phenotype.

**Figure 5.**
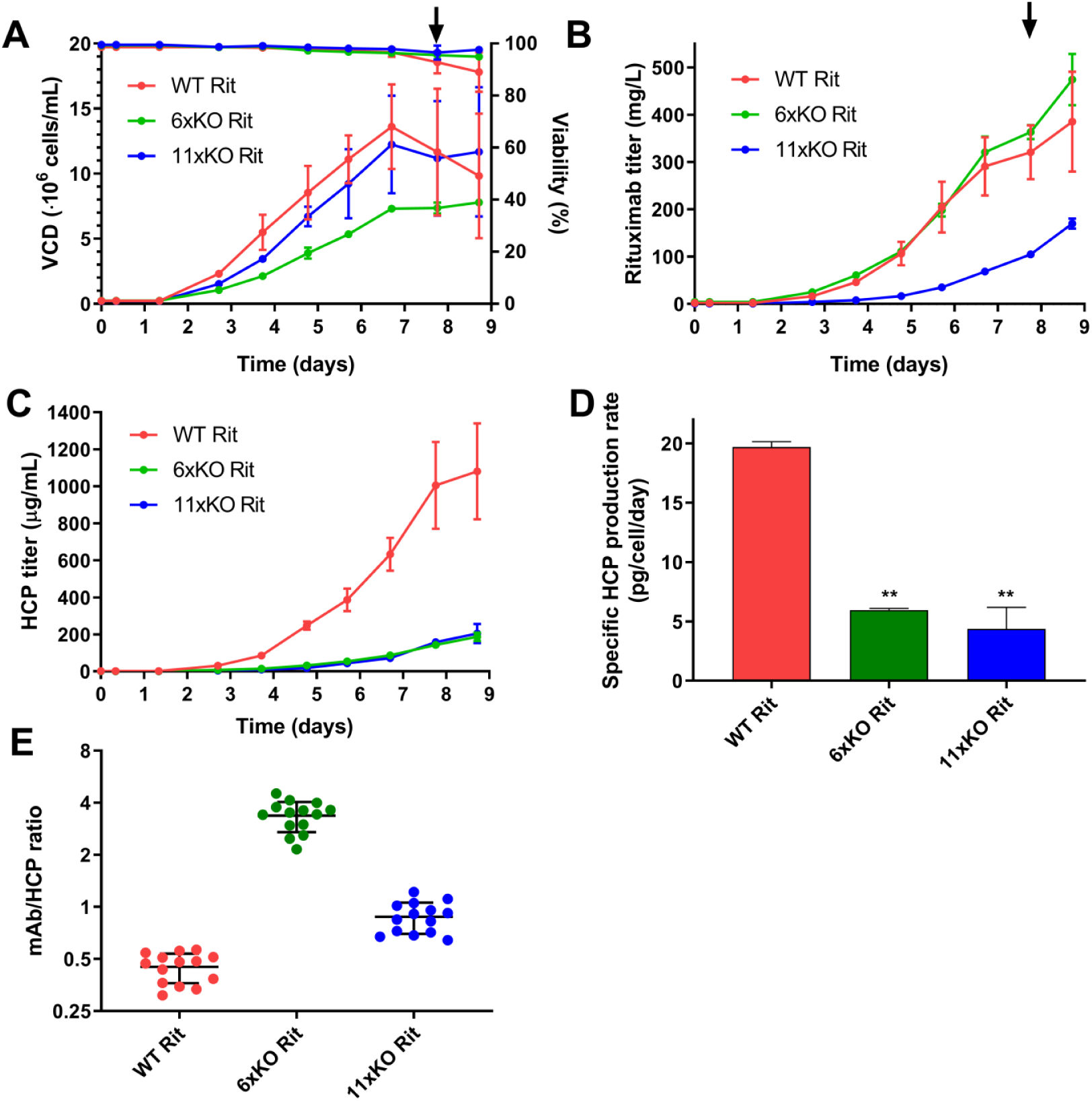
Antibody-producing clones retain the HCP-reduced phenotype under fed-batch cultivation. Viability and VCD (A), Rituximab titer (B), HCP profile (C), specific HCP productivity (D) and the mAb/HCP ratio (E) of WT Rit (red), 11×KO Rit (green), and 11×KO Rit (blue) cell lines. Viability, VCD, titer and HCP content were measured daily in a DASGIP bioreactor. At the indicated time point (arrow), 100 mL of the culture was harvested for mAb purification. The HCP/mAb ratio was calculated by dividing the mAb titer by the HCP titer for each time point. Mean and standard deviation of two technical replicates are shown. Statistical analysis was performed using one-way ANOVA followed by Tukey’s post-hoc test (** P ≤ 0.01).

Media screening using the Ambr bioreactor revealed that high-producing mAb clones could be selected from all knockout mutants (Fig. S3). As productivity was most comparable across mutants in FortiCHO medium, we selected this medium to scale up cultivation in a DASGIP bioreactor (Fig. 5). To limit the contribution of lysed cells to HCP content in the downstream process, we harvested when the first culture dropped below 90% viability (Fig. 5A, arrow). Maximum VCD of WT Rit and the 11×KO Rit cell lines were comparable (Fig. 5A), while the VCD of the 11×KO Rit cell line was approximately 50%lower (also observed in the Ambr bioreactor). However, Rituximab titer at harvest (Fig. 5B) of the 11×KO Rit cell line was comparable to WT Rit, while the 11×KO Rit cell line produced approximately 40% less. HCP titer reduction of the 11×KO Rit and 11×KO Rit cell lines was even more pronounced under these circumstances. Whereas the WT Rit cell line produces 1080 µg/mL HCP, the 11×KO and 11×KO cell lines behave similarly and produce approximately 200 µg/mL HCP (Fig. 5C). Specific HCP productivity was 19.7 pcd for WT Rit, whereas the 11×KO Rit and 11×KO Rit cell lines were reduced to 5.9 and 4.8 pcd, respectively (Fig. 5D). To emphasize the substantial improvement in product purity, we calculated the mAb/HCP ratio, which is improved 7-fold in the 11×KO Rit cell line and 2-fold in the 11×KO cell line (Fig. 5E). Importantly, under controlled cultivation, our knockout cell lines retain the strongly HCP-reduced phenotype.

### Knockout cell lines show reduced HCP content throughout a mAb purification process

To assess the effect of the knockouts on downstream purification, we performed protein A affinity chromatography, followed by cation exchange chromatography (CIEX) and anion exchange chromatography (AIEX) (see Fig. S4). Samples were collected after every chromatographic step and were analyzed with respect to mAb concentration and HCP content. Using these data, we calculated mAb purification efficiency, HCP content per mg mAb in ppm, and HCP log reduction values (LRV) (Table 2). Purification efficiency ranges from 35% to 55% and total HCP LRV of the process ranges from 4.4 to 5.8. HCP content after three chromatographic steps was 45 ppm, 18 ppm and 3 ppm for the WT Rit, 11×KO cell and 11×KO cell lines, respectively. In addition to the decrease in HCP content in the 11×KO cell line throughout mAb purification, we also observed an increase in HCP clearance during protein A chromatography of one order of magnitude. We conclude that the HCP-reduced phenotype is maintained throughout a representative downstream process.

**Table 2.**
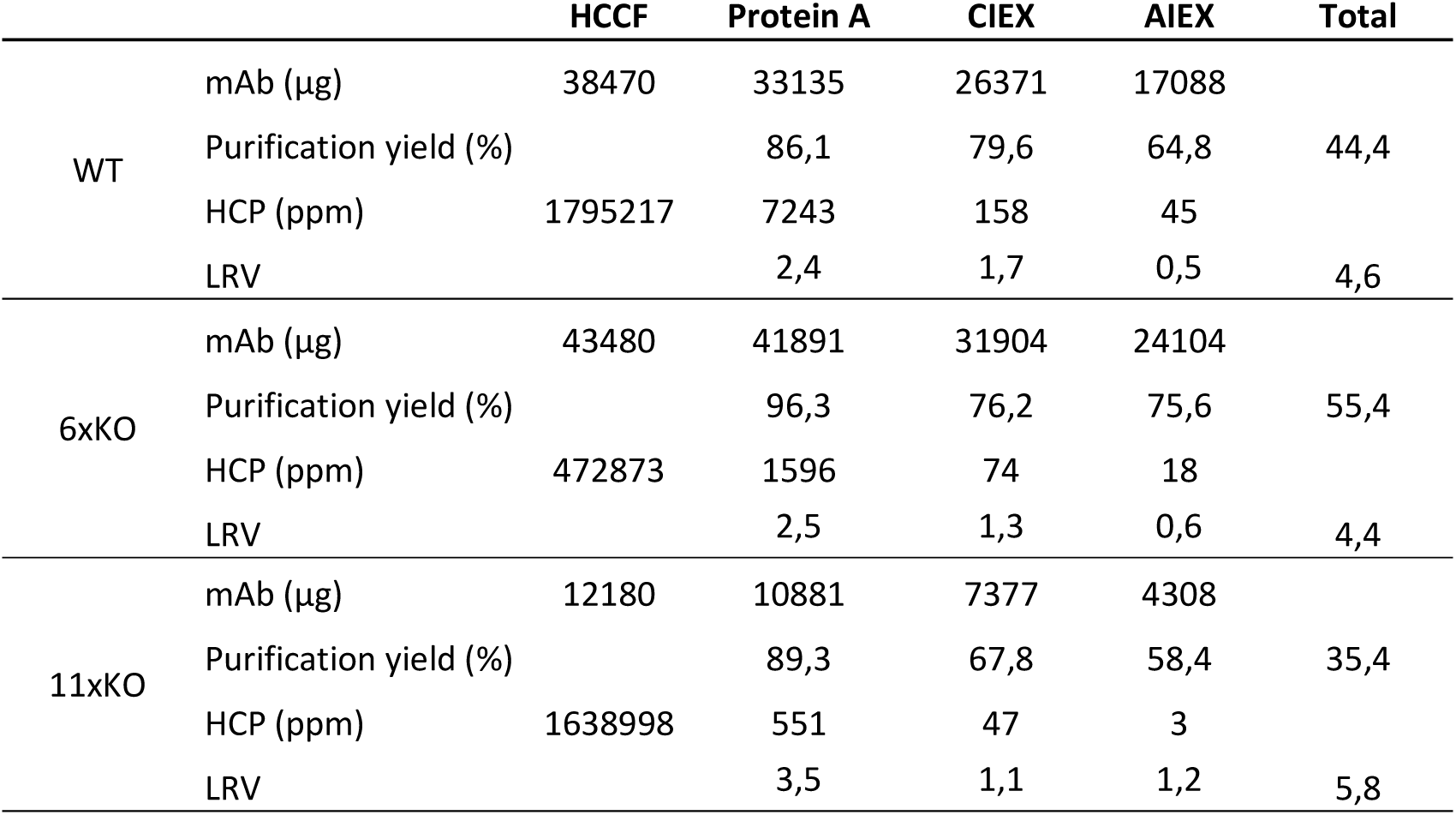
Knockout cell lines show reduced HCP content throughout a three-step purification process. The HCCF, protein A elution pool, cation exchange (CIEX) elution pool, and anion exchange (AIEX) flow through pool of WT Rit, 11×KO Rit and 11×KO Rit were subjected to mAb and HCP titer measurements. These values were used to calculate yield, HCP log reduction values (LRV), and HCP content in parts per million (ppm). Mean of technical duplicates is shown.

|  |  | HCCF | Protein A | CIEX | AIEX | Total |
| --- | --- | --- | --- | --- | --- | --- |
| WT | mAb (µg) | 38470 | 33135 | 26371 | 17088 |  |
|  | Purification yield (%) |  | 86,1 | 79,6 | 64,8 | 44,4 |
|  | HCP (ppm) | 1795217 | 7243 | 158 | 45 |  |
|  | LRV |  | 2,4 | 1,7 | 0,5 | 4,6 |
| 6xKO | mAb (µg) | 43480 | 41891 | 31904 | 24104 |  |
|  | Purification yield (%) |  | 96,3 | 76,2 | 75,6 | 55,4 |
|  | HCP (ppm) | 472873 | 1596 | 74 | 18 |  |
|  | LRV |  | 2,5 | 1,3 | 0,6 | 4,4 |
| 11xKO | mAb (µg) | 12180 | 10881 | 7377 | 4308 |  |
|  | Purification yield (%) |  | 89,3 | 67,8 | 58,4 | 35,4 |
|  | HCP (ppm) | 1638998 | 551 | 47 | 3 |  |
|  | LRV |  | 3,5 | 1,1 | 1,2 | 5,8 |

## Discussion

The yield of recombinant proteins during biomanufacturing has steadily increased over the past decades. Thus, downstream processing has become a bottleneck in biotherapeutic production^29^. Efforts to alleviate this have mostly focused on innovations in process design to improve downstream processing capacity, speed, and economics^30,31^. It has also been suggested that cell line engineering could be used to reduce or remove problematic HCPs, lessening the need for additional purification steps^6,32^. Indeed, researchers have specifically removed unwanted proteins from CHO cell lines that affect product quality^1,2,11,12^. In addition, it has recently been shown that depletion of a highly abundant mRNA encoding a non-essential protein can improve growth and product titer, indicating that it is possible to free up energetic and secretory resources by rational cell line engineering strategies^33^. This observation was supported by a model of the secretory pathway, which was able to accurately predict the increase in growth and protein productivity^19^. These studies have so far been limited to single engineering targets, but could easily be applied on a larger scale. Although different in approach, a recent paper pioneers the use of inducible downregulation of non-product related protein expression leading to an increase in the purity of the biopharmaceutical product^34^. Knowledge-based decisions can support purity-by-design approaches and could lead to easier downstream purification processing.

Here, we hypothesized that removal of HCPs by targeted gene disruption will lead to improvements in cell growth, secretory pathway capacity, and DSP. We used computational modeling to demonstrate that CHO cells spend substantial resources on the host cell secretome. We proceeded with the removal of 14 HCPs, which were more highly abundant in CHO HCCF. We also prioritized targets that can be difficult-to-remove or that influence product quality. Surprisingly we found the removal of these 14 HCPs led to a substantial decrease in HCP content of up to 70% and HCCF total protein content of up to 50%. The observed phenotype persists under controlled cultivation and is specifically caused by the knockouts. Importantly, the ability to generate high mAb producers was not perturbed.

Improvements in product purity will considerably facilitate the purification of mAbs to acceptable HCP levels, especially in those situations where HCPs form a challenge in the purification or quality of a biotherapeutic protein. It is tempting to speculate that one of the ion exchange chromatography steps could be omitted from the downstream purification process. However, besides the removal of HCP impurities, ion exchange chromatography is also used to reduce high molecular weight aggregates, charge-variants, residual DNA, leached Protein A and viral particles^35^. It remains to be seen whether two chromatographic steps can bring other product- and process-related impurities within regulatory requirements. The results presented here certainly warrant a detailed analysis of the HCP-reduced cell lines under industrial mAb production conditions with respect to growth, production and mAb critical quality attributes.

## Conclusion

Through a multiplex genome editing approach here, we successfully removed more than half of the HCPs in CHO cells by mass. This demonstration highlights that large-scale genome editing in mammalian cells can be done for bioproduction, and can have benefits throughout the bioproduction process. In particular, such efforts can be effectively used to make clean CHO cells by eliminating many undesirable HCPs^6^ and retroviral-like particles^36^ that currently require extensive purification strategies. Through this, one may obtain higher quality drugs and produce these at lower costs.

## Author contributions

S.K., D.L., H.F.K., G.M.L. and N.E.L. designed the experiments. SK, T.W. and L.E.P. selected knockout targets. T.W. performed proteomic analysis. S.K., M.D. and D.L. performed cell line cultivation, selection and analysis. S.K. performed protein purification and analysis. N.E.L., J.M.G. and A.W.T.C. performed *in silico* analysis. S.K. and N.E.L. wrote and corrected the manuscript.

## Acknowledgments

The authors would like to thank Bjørn Gunnar Voldborg, Helle Munck Petersen, Christian Oscar Wistrøm, Mikkel Schubert, Zulfiya Sukhova, Karen Kathrine Brøndum, Elham Maria Javidi, and Kristian Lund Jensen for support. This work was supported in part by the Novo Nordisk Foundation (NNF10CC1016517). N.E.L. and J.M.G. acknowledges support from NIGMS (R35 GM119850), and a fellowship from the Government of Mexico (CONACYT) and the University of California Institute for Mexico and the United States (UC-MEXUS).

## Supplementary material

**Table S1. Proteomic analysis of WT CHO-S HCPs.** UniProt identifiers, protein names and relative abundance of the 200 most abundant proteins found in a CHO-S supernatant are shown. The targets addressed in this study are indicated in bold. Abundance values are the mean of technical triplicates.

See Suppl Table S1 Kol et al.

### Text to figure S1

A possible explanation for the HCP-reduced phenotypes observed in this study is that the creation of knockouts selects for faster-growing cells. As cells have go through a single-cell stage where survival rates are low, a selective pressure for better growth is imposed. It may thus be possible that the phenotype is not caused by specific knockouts, but is a more general effect of global downregulation of costly, secretory proteins. We quantified the HCP content of cell lines generated in an independent study addressing glycosylation^1^. These cell lines were subjected to 3 cycles of CRISPR/Cas9-mediated multiplex gene disruption. In the first round, 6 genes were disrupted. In the second round, two genes were disrupted. Finally, in the third round, another two genes were disrupted, resulting in a cell line harbouring 10 gene disruptions. No significant decrease in HCP content of WT CHO-S and the knockout cell lines is observed showing that the HCP-reduced phenotype is specifically caused by the removal of highly abundant proteins.

**Figure S1.**
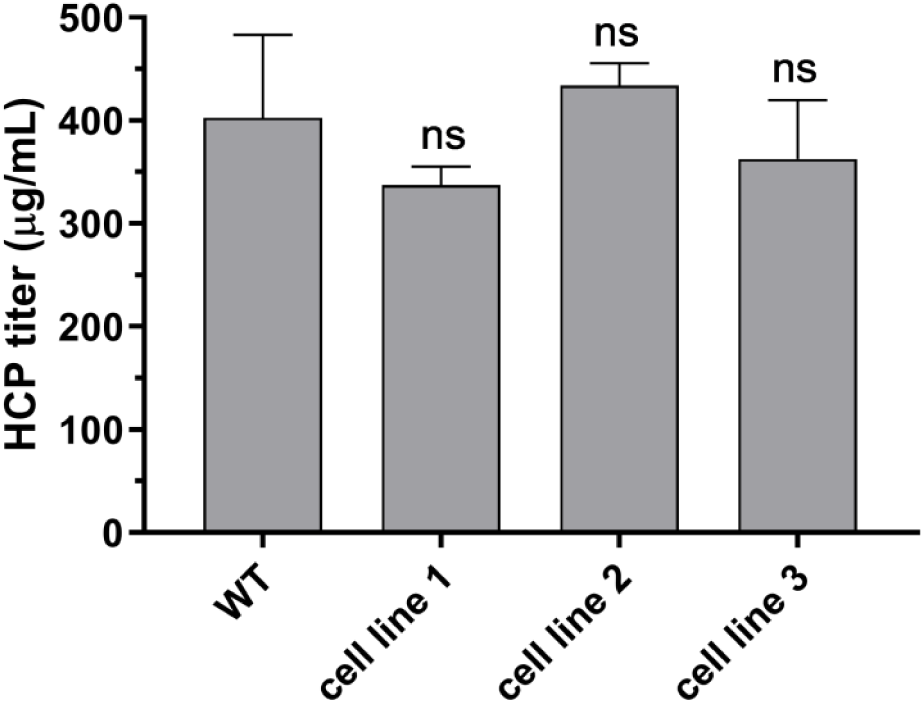
HCP content of WT CHO-S and cell lines harbouring multiple gene disruptions. HCP content was measured of WT CHO-S (WT) and cell lines harbouring 6 gene disruptions (cell line 1), 8 gene disruptions (cell line 2), and 10 gene disruptions (cell line 3). HCP content was measured after cultivation for four days in shake flasks. Mean and standard deviation of three technical replicates are shown. Statistical analysis was performed using one-way ANOVA followed by Tukey’s post-hoc test (ns = not significant).

**Figure S2.**
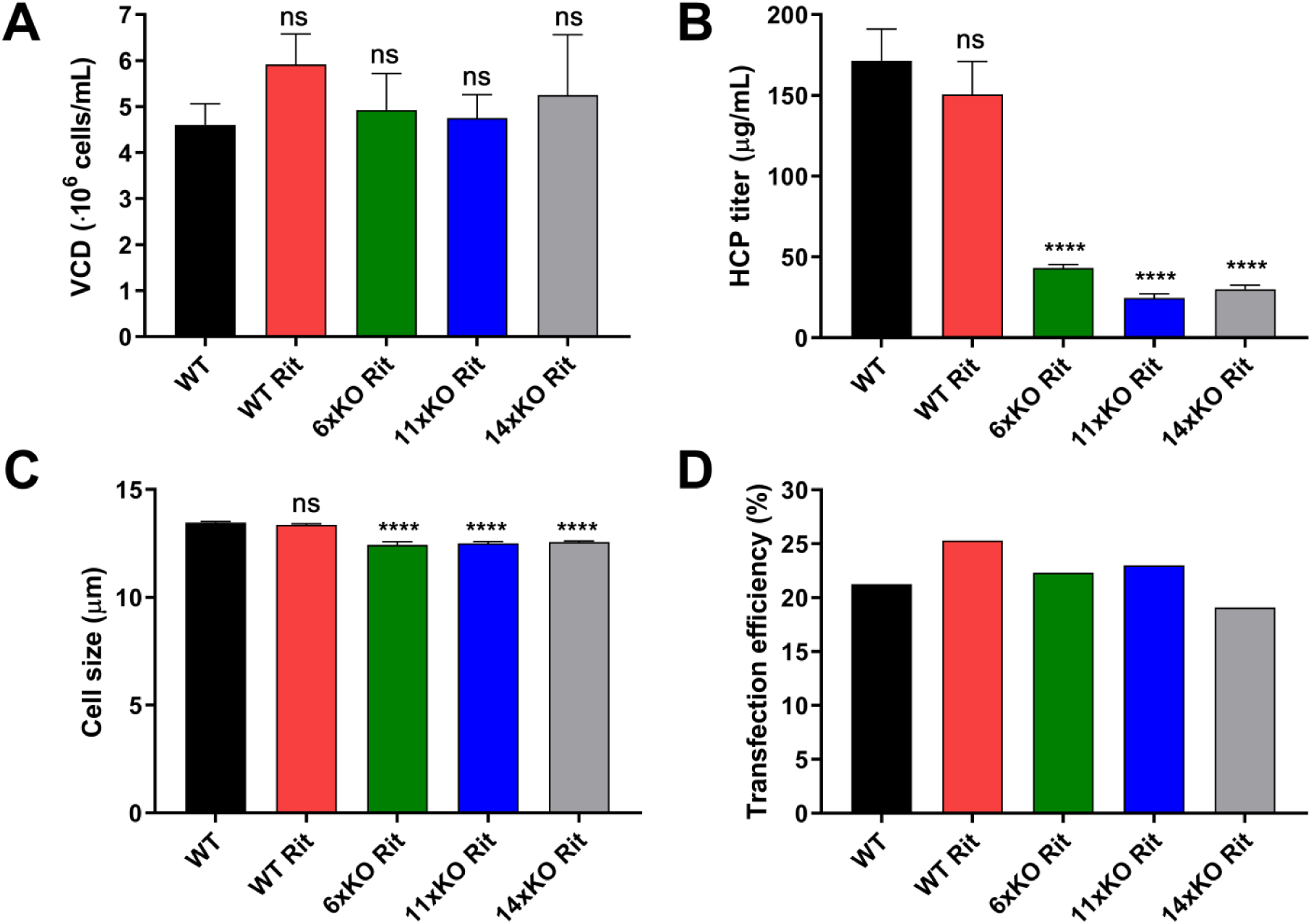
VCD, HCP content and cell size. VCD (A), HCP content (B), cell size (C) and transfection efficiency (D) of WT (black), WT Rit (red), 11×KO Rit (green), 11×KO Rit (blue), and 11×KO Rit (grey) cell lines. All properties were measured after cultivation for four days in shake flasks. Mean and standard deviation of three technical replicates are shown for all experiments except for the transfection efficiency determination. Statistical analysis was performed using one-way ANOVA followed by Tukey’s post-hoc test (ns = not significant, **** P ≤ 0.0001).

## Text to figure S3

Growth characteristics and productivity of the knockout mutants were evaluated in an Ambr bioreactor using FortiCHO, ActiPro and OptiCHO media. Productivity was found to be highest in the 11×KO Rit cell line in OptiCHO medium (Fig. S3C, right panel). VCD of all clones cultured in this medium reached a maximum level of approximately 15 million cells per milliliter. While the VCD of WT Rit and 11×KO dropped after 8 days in culture, VCD of 11×KO Rit was constant for the remainder of the experiment (Fig. S3A, right panel). In addition, viability of the 11×KO Rit cell line remained higher than the other cell lines in this medium (Fig. S3B, right panel), indicating prolonged culture longevity. In ActiPro medium, we observed low VCD and productivity of both knockout cell lines as compared to WT Rit (Fig. S3A and S3C, center panels). In FortiCHO medium, the productivity of all cell lines was comparable, even though the cell density of WT Rit was higher than the 11×KO Rit and 11×KO Rit cell lines (Fig. S3A and S3C, left panels). Considerable differences in cell density, viability and titer were observed depending on the culture medium. These data show that high mAb production can be achieved in our knockout cell lines.

**Figure S3.**
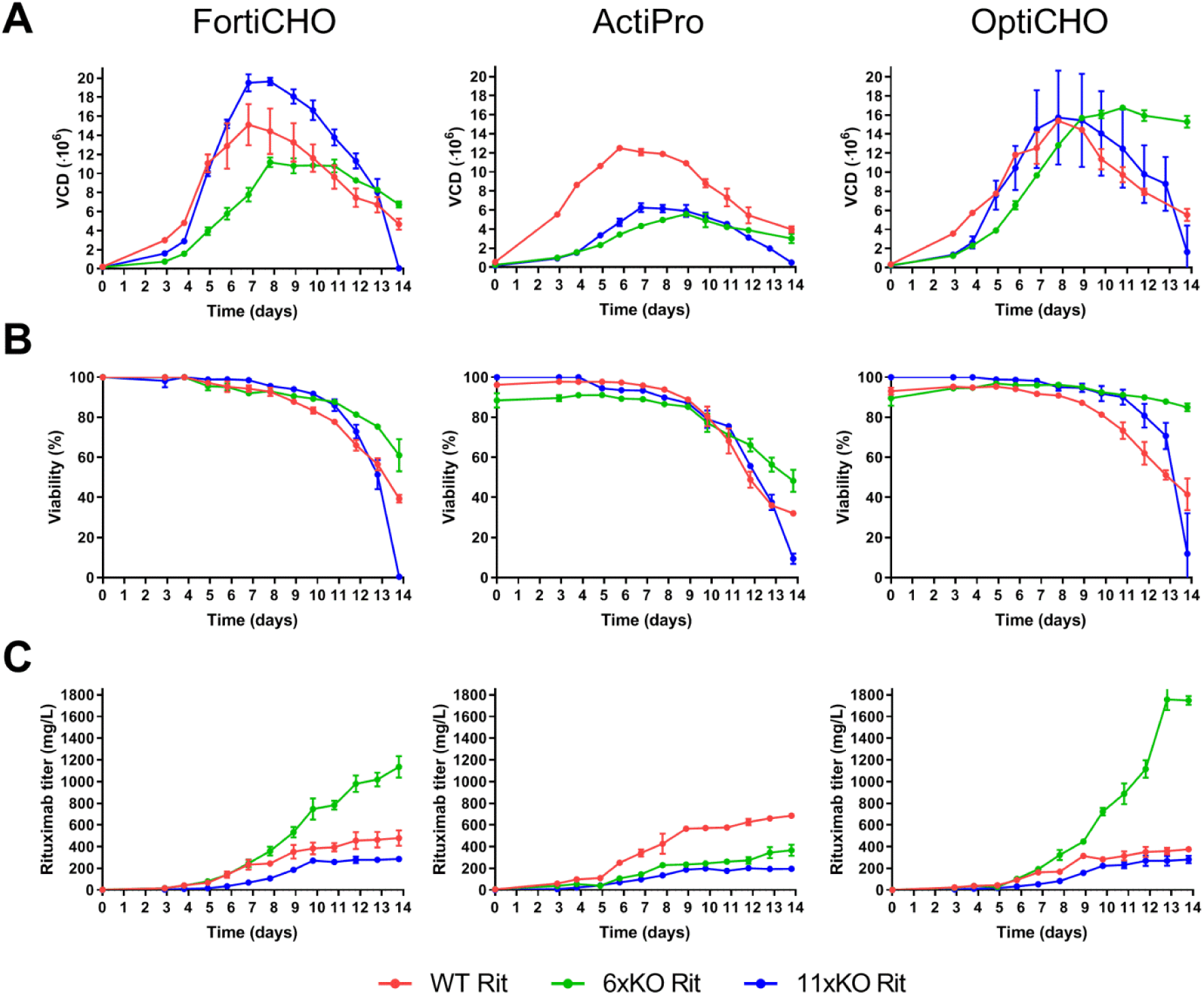
Growth characteristics and productivity of the cell lines described in this study under fed-batch conditions in an Ambr bioreactor. VCD (A), viability (B), and titer (C) of WT Rit (red), 11×KO Rit (green), and 11×KO Rit (blue) measured daily during cultivation in an Ambr bioreactor. Cell lines were cultured in FortiCHO (left panels), ActiPro (center panels), and OptiCHO media (right panels). Mean and standard deviation of three technical replicates are shown.

**Figure S4.**
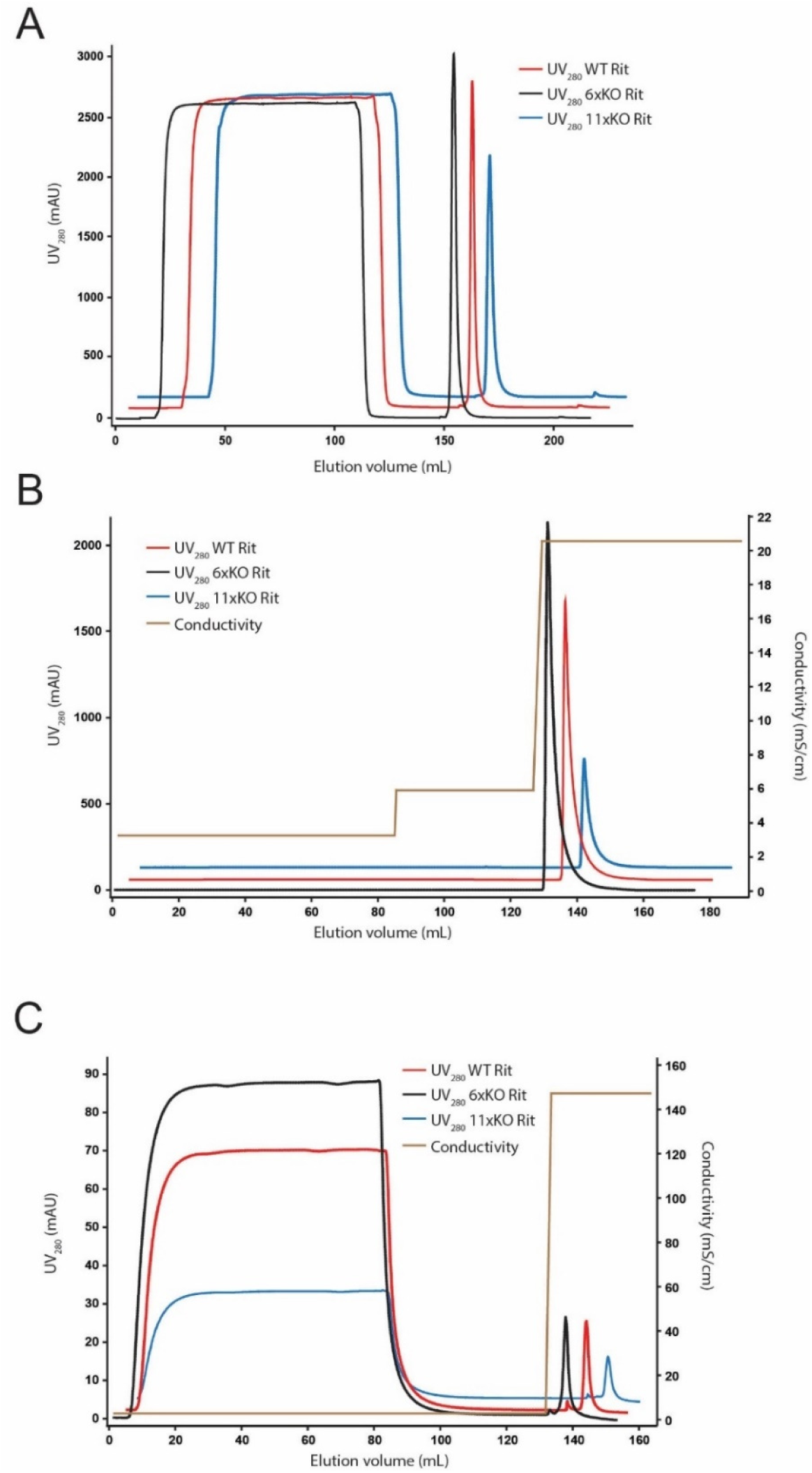
Protein purification chromatograms. Representative protein A (A), CIEX (B) and AIEX (C) chromatograms of the purification of Rituximab produced in WT Rit (red line), 11×KO Rit (green line) and 11×KO Rit (blue line). Protein A and C3IEX chromatography were run in bind-and-elute mode, while AIEX chromatography was run in flow-through mode. The conductivity profile during CIEX and AIEX is indicated (brown line).

